# InSituPy – A framework for histology-guided, multi-sample analysis of single-cell spatial transcriptomics data

**DOI:** 10.1101/2025.03.07.641860

**Authors:** Johannes Wirth, Anna Chernysheva, Birthe Lemke, Isabel Giray, Aitana Egea Lavandera, Katja Steiger

**Affiliations:** Institute of Pathology, School of Medicine and Health, Technical University Munich, Munich, Germany

**Keywords:** Spatial transcriptomics, computational framework, in situ sequencing

## Abstract

Single-cell spatial transcriptomics (scST) methodologies allow, in combination with histological stainings, an unprecedented view on disease progression. To comprehensively analyze scST data, bioinformatic analysis frameworks need to integrate the diverse set of data modalities and, just as importantly, enable the joint analysis of multiple datasets from clinical or experimental cohorts together with its corresponding metadata. Here, we present the InSituPy framework to comprehensively analyze single-cell spatial transcriptomic data from a multi-sample level down to the cellular and subcellular level. The framework contains analysis workflows for the integration of image data as well as pathological and biological expert knowledge. Increasing the accessibility of the data for non-bioinformaticians, the framework opens new ways of generating hypotheses, especially in the context of translational research.

## Introduction

The emergence of novel multi-omics technologies has revolutionized our understanding of biological processes in healthy and diseased tissue. Single-cell transcriptomics methods allow researchers to investigate the cellular composition of tissues but, because a dissociation of the cells is required, lead to a loss of the spatial context. Spatially resolved transcriptomics (ST) technologies overcome this limitation and enable researchers to investigate the gene expression of cells while preserving the information about the spatial location within the tissue. While early methods such as Visium and Slide-seq failed to achieve single cell resolution, the development of multiplexed fluorescence in situ hybridization (multiplexed FISH)^1,2^ and in situ sequencing (ISS)^3,4^ allowed researchers to map individual RNA molecules at subcellular resolution and thus measure the transcriptional state of single cells of a tissue sections. The non-destructive nature of multiplexed FISH and ISS technologies allows the combination of transcriptomic readouts with conventional image-based readouts such as histological or immunofluorescence stainings. The access to histological data makes it possible for the first time to fully integrate knowledge of pathology experts from routine diagnostics and creates a new way of forming hypothesis. Further, recent technologies such as Xenium in Situ and MERSCOPE allow the analysis of multiple tissue sections at once, increasing the throughput of the methods and enabling the generation of large clinical cohorts, e.g. using tissue microarrays. To fully exploit the potential of such datasets, it is important to integrate experiment-level information (e.g. clinical metadata or treatment conditions) with the single-cell spatial transcriptomics (scST) data. While Python-based frameworks such as SpatialData^5^ allow an integration of ISS data modalities, they do not offer a way to structurally integrate data from multiple tissue sections and corresponding metadata. Additionally, since a growing number of researchers uses scST technologies in their projects, an easy applicability, also for non-bioinformaticians, is key. In this publication, we present InSituPy, a framework to explore and analyze ISS data while simplifying and accelerating the preprocessing of multiple large datasets in parallel. Using this framework we introduce workflows to integrate pathological expert knowledge into the analysis.

## Results

### Overall structure of the framework

Single-cell spatial omics datasets consist of multiple data levels (Figure 1 A). These levels comprise (i) the experiment level, which can contain information about the patients, the model system or treatments; (ii) the sample level, containing all data of one data entity, e.g. one tissue section; (iii) the cellular level with the method specific readout per single cell, e.g. gene or protein expression; and (iv) the subcellular level including information such as individual transcripts. InSituPy offers a data structure to read, analyze and store data from all those four levels, while structuring them in a biologically meaningful manner (**Figure 1 B** and **Supplementary Figure 1**). Divided into two main data objects, *InSituExperiment* and *InSituData*, the framework facilitates the integration of data from multiple tissue sections per experiment across multiple experiments (**Supplementary Figure 1 B**). An InSituExperiment object consists of multiple InSituData objects, with each InSituData object containing the scST data from one sample (Figure 1 C). Within the InSituExperiment object, each InSituData object gets assigned a unique ID (UID) to connect the datasets with its corresponding metadata, making it possible to easily query datasets based on their metadata, integrate experimental-level metadata into the analysis, iterate through multiple samples or concatenate data from multiple experiments (**Supplementary Figure 1 D-G**). Importantly, the structure of InSituPy is designed to be compatible with any spatially resolved single-cell transcriptomics data consisting of a gene/cell count matrix, image data and optionally segmentation masks or transcript locations. Currently, only readers for the Xenium In Situ technology are available, but the open-source publication of the code and a generalized data structure will facilitate the implementation of community-developed data readers for other technologies in the future. InSituPy is tested on all common operating systems such as Windows, Linux and Mac.

**Figure 1.**
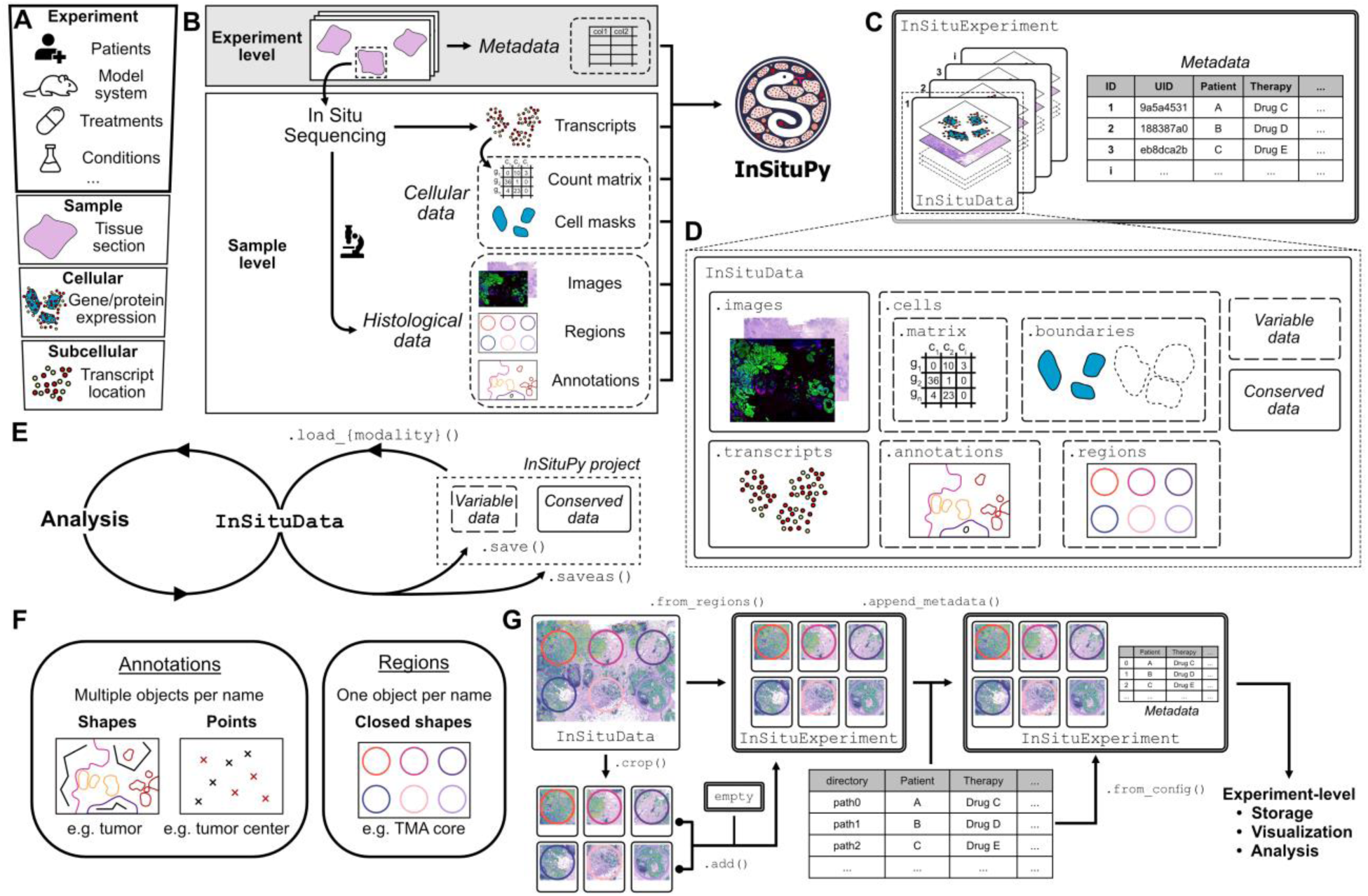
Overview of InSituPy package structure. **(A)** Overview of data levels available in state-of-the art single-cell spatial transcriptomics datasets. **(B)** Schematic illustrating the experiment-level and sample-level data acquisition steps of in situ sequencing (ISS) experiments and the data integration into the InSituPy framework. **(C)** and **(D)** Schematics showing the data structure of the InSituExperiment class and the InSituData class, respectively. An index and a unique ID (UID) connect the datasets and the metadata within the InSituExperiment class. In an InSituData object, data is structured into *images*, *cells*, *transcripts*, *annotations* and *regions*, where cells is further subdivided into the transcriptomic data (*matrix*) and the data of the cellular *boundaries*. Further, the data is divided into variable and conserved data, where variable data is updated during the analysis and conserved data stays constant as shown in **E**. **(F)** Illustration to show the differences between histological annotations and regions. **(G)** Schematic showing different possibilities to generate an InSituExperiment object.

### Data structure on sample level

On the sample level, i.e. within an InSituData object, data is organized in five data layers: cellular data (“.cells”), aligned image data (“.images”), transcript locations and identities (“.transcripts”), histological annotations (“.annotations”), and regional annotations (“.regions”) (Figure 1 D). The cellular data in turn consists of two sublayers: the single-cell transcriptomics data (“.matrix”) and the cellular boundaries (“.boundaries”). To save memory, all data modalities can be loaded individually as required in the analysis using the respective loading functions (**Supplementary Figure 1 A**). During analysis with InSituPy, data can be saved in an InSituPy project folder. When data is saved to an existing InSituPy project, the framework differentiates between conserved and variable data. Conserved data comprises all data that does not change during analysis (i.e. image data and transcript locations), while variable data is data that is modified during the analysis, i.e. the single-cell transcriptomics data, the cellular boundaries as well as the regional and histological annotations (**Figure 1 E**). When saving an existing InSituPy project, only the variable data is changed on disk while the conserved data remains unchanged. This significantly accelerates saving processes and reduces the amount of storage space required (**Supplementary Figure 2**).

### Data formats

The framework uses different state-of-the-art packages for data handling. An overview of all modalities and respective data formats can be found in **Supplementary Table 1**. Single-cell transcriptomics data is loaded in the AnnData format and stored as *h5ad* file. To efficiently load large image data, InSituPy exploits the resource-saving capabilities of the *dask* framework and stores the data in the *Zarr* format allowing lazy loading of the data. Alternatively, to achieve easy compatibility with common image analysis and annotation tools such as QuPath, the image data can be also saved in the widely used OME-TIFF format. Histological data such as geometric shapes are processed using the *GeoPandas* framework and saved as GeoJSON file.

### Classification of histological data

In case of histological data, InSituPy differentiates between regions and annotations (Figure 1 F). Annotations consist of any kind of geometric shapes (e.g. polygons, lines or points) with each shape getting assigned to a certain *class* (e.g. “tumor cells”, “immune cells”) and a *key* (e.g. the name of the pathologist doing the annotations). In case of annotations, within one key multiple shapes are allowed to have the same name (multiple annotations could be named “tumor cells”) and a unique identifier is used to differentiate between the polygons. Regions in turn can only consist of polygons and are required to have a unique class name, i.e. only one polygon per class name is allowed. Regions can delineate the positions of TMA cores or the position of different tissue sections or regions of interests within the same dataset.

### Multi-sample analysis using InSituExperiment

Current bioinformatic frameworks for the analysis of spatial transcriptomics data do not capture the experiment level which usually includes multiple datasets. InSituPy enables the integration of multiple datasets using its InSituExperiment class (**Figure 1 C**). Such a multi-sample dataset can be assembled using different strategies (**Figure 1 G**): (i) histological regions can be used to generate it, e.g. if one dataset contains multiple tissue sections or TMA cores, each originating from different samples. (ii) Individual InSituData object can be added manual to an existing InSituExperiment. Afterwards, sample-specific metadata can be assigned to the data (Figure 1 G). And (iii), an InSituExperiment can also be created based on a configuration file containing the data directories and the corresponding metadata. Based on the InSituExperiment class, InSituPy allows the storage, visualization and analysis of multi-sample scST datasets. This includes functions to query the metadata and return the resulting datasets, iterate through datasets, apply functions to all datasets, concatenate multiple scST datasets, or show an overview of QC metrices across all datasets (**Supplementary Figure 1 D-H**). Compared to other state-of-the-art frameworks for spatial transcriptomics data, InSituPy is the only which allows such a multi-sample analysis while also offering a comprehensive analysis on the individual sample level (**Supplementary Table 2**).

### Automated image registration

The non-destructive approach of scST methods such as Xenium In Situ enable the subsequent staining of sections using immunofluorescence (IF), immunohistochemistry (IHC) or hematoxylin and eosin (H&E). To enable pathological annotations, this image data needs to be aligned with the scST data. InSituPy contains an automated registration pipeline to simplify this task and align the subsequently stained images to the nuclear image acquired during the scST measurement (**Figure 2 A**). *Scale-Invariant Feature Transform* (SIFT)^6^ and the *Fast Library for Approximate Nearest Neighbors* (FLANN)^7^ are used to identify and match features in both images. If not enough matched features are found, the algorithm automatically tests whether flipping the image vertically or horizontally leads to better results and continues with the best of these options. Subsequently, the *random sample consensus algorithm* (RANSAC) is used to select the most robust matches and calculate an affine or perspective transformation matrix for registering the images. After registration of the images, they are saved together with QC images and can be automatically loaded using InSituPy.

**Figure 2.**
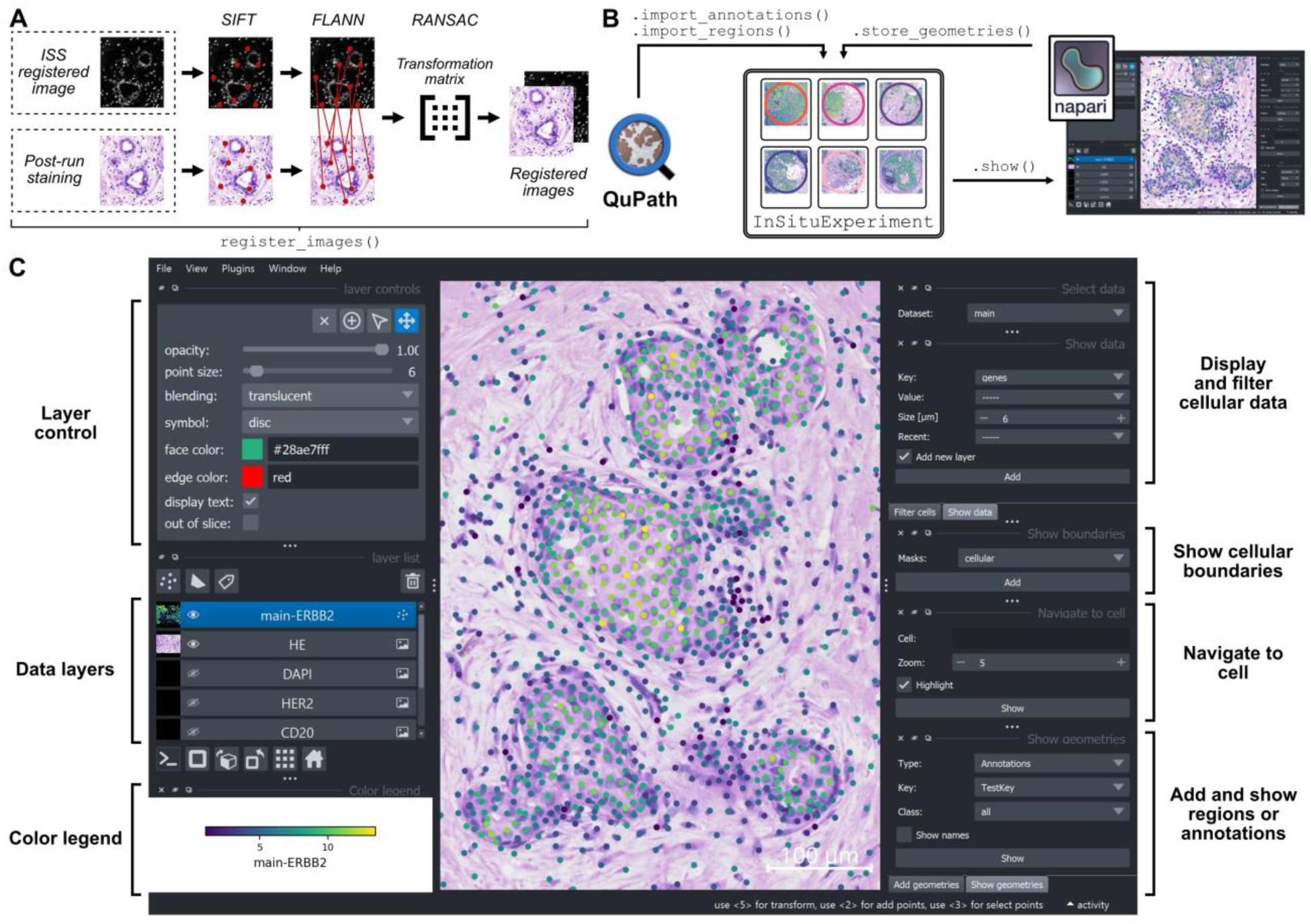
Automatic image registration, annotation and visualization of data in InSituPy. **(A)** Workflow showing the steps for automatic image registration of images from post-run stainings with a fluorescent image that is already registered to the transcriptomic data. *Scale-Invariant Feature Transform* (SIFT)^6^ and the *Fast Library for Approximate Nearest Neighbors* (FLANN)^7^ are used to identify and match features in the two images. The *random sample consensus algorithm* (RANSAC) is used to select the most robust matches and calculate an affine or perspective transformation matrix for registering the images. **(B)** Schematic showing the possible ways to import geometries (annotations or regions) into InSituPy, either from external tools such as QuPath or from the napari based data viewer. **(C)** Screenshot showing the interactive napari^8^ viewer integrated in InSituPy.

### Data visualization and annotation using the InSituPy viewer

A precise registration of the images is a prerequisite for the addition of histological annotations to the datasets and for visualization of the scST cell type annotations for pathologists. The latter especially being important in terms of morphological quality control, but also to leverage the potential of scST analyses in a translational setting. Histological annotations can be either imported from external software such as QuPath^9^ or from a napari-based^8^ viewer (**Figure 2 B**). Within the viewer, different functionalities are implemented, that allow the exploration of the data, including functionalities to visualize transcriptomic data, cellular boundaries, locating specific cells as well as the addition and display of annotations or regions (**Figure 2 C** and **Supplementary Figure 3**). Subsequently, added regions or annotations can be stored within the source InSituExperiment object or, alternatively, they can also be imported from external software (e.g. QuPath^9^) using the corresponding functions (Figure 2 B).

### Experiment-level analyses using InSituPy

Preprocessing of the data, including normalization and transformation as well as dimensionality reduction and clustering, are central steps in the analysis of in situ sequencing data. InSituPy contains simplified workflows to apply such steps across multiple datasets in a reproducible manner (**Supplementary Figure 1 C**). Further, the framework contains functions to conduct histology-guided analyses on an experiment level. Utilizing the structure of the InSituExperiment object, InSituPy enables an analysis within samples or between different samples and depending on the biological question, the analysis can be focused on specific samples, annotations or cell types (**Figure 3 A**). The currently available analyses include the evaluation of the cellular composition, the measurement of cell type densities, differential gene expression analysis and GO term enrichment analysis (**Figure 3 A)**. For the latter, InSituPy offers interfaces to STRING^12^, Enrichr^14^ and g:profiler^13^ and provides plotting functionalities to visualize the results. Together, these functionalities allow a histology and pathology-informed analysis of gene expression changes in multi-sample scST datasets.

**Figure 3.**
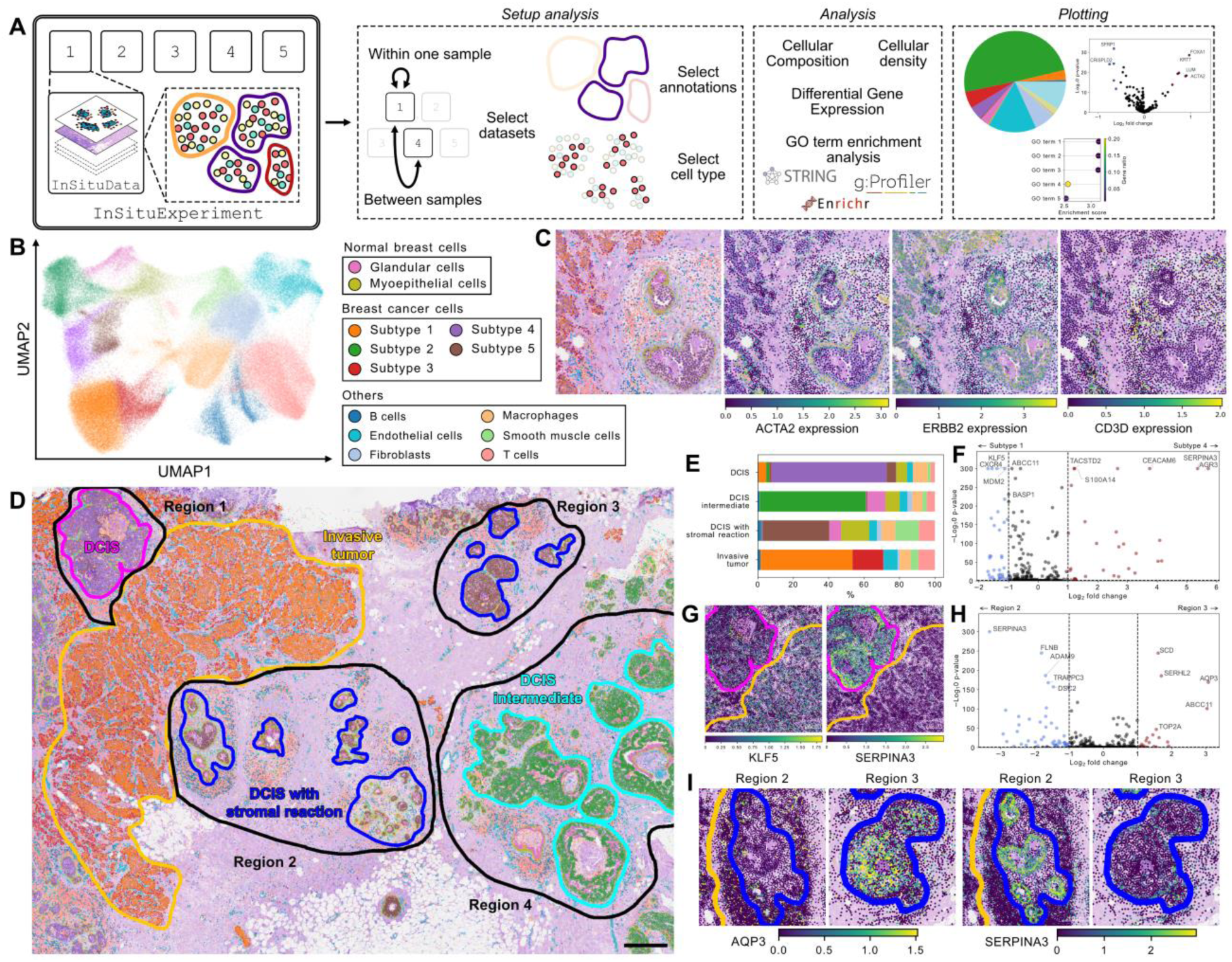
Histology-guided, multi-sample analyses using InSituPy. **(A)** Overview of the steps required to perform cross-sample analyses in InSituPy. The example dataset consists of five InSituData objects saved within an InSituExperiment class. First, the analysis setup needs to be specified, including the selection of the samples of interest, the annotations and the cell type. Subsequently, the type of the analysis needs to be chosen. Currently, following analyses are implemented: Cellular composition within annotations, differential gene expression (DGE) analysis, and GO term enrichment analysis using different upstream pipelines^12–14^. Also plotting of the results is implemented. **(B)** Two-dimensional UMAP embedding of an example breast cancer *Xenium In Situ* dataset^11^. Colors denote the cell type annotation. **(C)** Overlay of the transcriptomic data and an image of the H&E stained tissue section. Colors denote the cell type or the gene expression of *ACTA2*, *ERBB2* or *CD3D*, respectively. **(D)** Overlay of cells depicted as points and pathological annotations and regions on an H&E stained breast cancer section. Colors of the points denote the cell type. Scale bar: 500 µm. **(E)** Bar chart showing the results of the cellular composition analysis in the pathological annotations depicted in D. Colors correspond to cell types shown in B. **(F)** Volcano plot showing the differentially expressed genes in breast cancer subtype 4 within the “DCIS” annotation compared to subtype 1 within the “Invasive” annotation. **(G)** Cellular gene expression of *KLF5* and *SERPINA3* in an example region including both annotations of DCIS and invasive tumor. Annotation colors correspond to D. **(H)** Volcano plot showing the differentially expressed genes in cells of breast cancer subtype 5 in DCIS with stromal reaction in region 3 compared to region 2. **(I)** Cellular gene expression of *AQP3* and *SERPINA3* in region 3 and 2 respectively. Annotation colors correspond to D. UMAP: Uniform Manifold Approximation and Projection; H&E: Hematoxylin and eosin.

### Identification of breast cancer subtypes using InSituPy

To exemplify these steps, we used a published breast cancer dataset^10,11^ which has been generated with the Xenium In Situ technology. Marker gene-based cell type annotation revealed normal breast cells such as glandular cells and myoepithelial cells, five different subtypes of breast cancer cells as well as other cell types including immune cells and stromal cells (**Figure 3 B**). InSituPy provides visualization functionalities to explore the distribution of cell types in the sample as well as the expression of genes of interest (**Figure 3 C**). Mapping all cell types on the whole specimen revealed a distinct spatial clustering of the different breast cancer subtypes (**Figure 3 D**). Annotation of the H&E image by a pathologist revealed one larger area with invasive tumor and multiple smaller areas with *ductal carcinoma in situ* (DCIS). The DCIS regions were further divided into classical DCIS, DCIS with stromal reaction and an intermediate DCIS type. These annotations correspond well with the annotations of DCIS #1 and DCIS #2 in the original publication^10^. Using InSituPy functionalities to investigate the cellular composition of the different annotations and regions, revealed a clear correspondence of the different breast cancer subtypes with the pathological annotations (**Figure 3 E** and **Supplementary Figure 4 A**). The classical DCIS matched well with breast cancer subtype 4 while DCIS with stromal reaction largely consisted of subtype 5. The intermediate typed DCIS predominantly consisted of breast cancer subtype 2 and the invasive tumor of subtypes 1 and 3, demonstrating the value of pathological annotations for cell typing.

### Histology-guided differential gene expression analysis across datasets

A central task in transcriptomic data analysis is the detection of differentially expressed genes. Using the above-mentioned structure of the InSituPy framework (**Figure 3 A**), the analysis can be limited to selected annotations or cell types and results can be visualized in volcano plots. Here, we demonstrated this functionality by investigating gene expression differences between cells of breast cancer subtype 4, corresponding to classical DCIS and cells of subtype 1, corresponding to invasive tumor cells. The analysis was limited to the respective pathological annotations and revealed 30 up- and 37 downregulated genes ( **Figure 3 F** and **Supplementary Figure 4 B**). The expression of two example genes, *KLF5* and *SERPINA3*, in a representative region is shown in **Figure 3 G**. Next, we used the DGE analysis functionalities of InSituPy to compare the gene expression of one cell type, breast cancer subtype 5, between region 2 and region 3. The analysis was restricted to cells within the pathological annotations “DCIS with stromal reaction” and resulted in 21 up- and 49-downregulated genes (**Figure 3** and **Supplementary Figure 4 C**). Two of these genes, AQP3 and SERPINA3, are shown in **Figure 3 I** and reveal clear gene expression differences in cells of breast cancer subtype 5 which could indicate further subtypes or cell states within it and underline the importance of the integration of pathological information into the analysis. GO term analysis of the upregulated genes yielded a significant enrichment of processes connected to vascular morphogenesis and VEGF-mediated proliferation signals (**Supplementary Figure 4 D**), which might be connected to the pronounced vascularization in close proximity of the tumor cells in region 3 (**Supplementary Figure 4 E**).

### Exploration of distance-dependent changes

The distance to neighboring cells or anatomical structures as well as the density of cells influences the phenotype of cells. Exploring such distance-dependent changes can provide information about pathological processes. Utilizing either kernel density estimation or the *mellon* package for cell-state density estimation in single-cell data^17^, InSituPy calculates the density of particular cell types within the samples, e.g. breast cancer cells or T cells (**Figure 4 A**). Further, InSituPy provides streamlined workflows to simplify such analyses, starting with the calculation of cell-cell distance relationships within the sample, as exemplified in **Figure 4 B** using the tumor centers as reference points. Subsequently, the cellular composition of the tumor can be visualized as a function of the distance to the tumor center, revealing distance-dependent changes in the tumor microenvironment (**Figure 4 C**). Further, InSituPy allows the exploration of distance-dependent expression changes of selected genes, as exemplified here for *ACTA2*, *LUM*, *MMP2*, and *CXCL12* (**Figure 4 D**).

**Figure 4.**
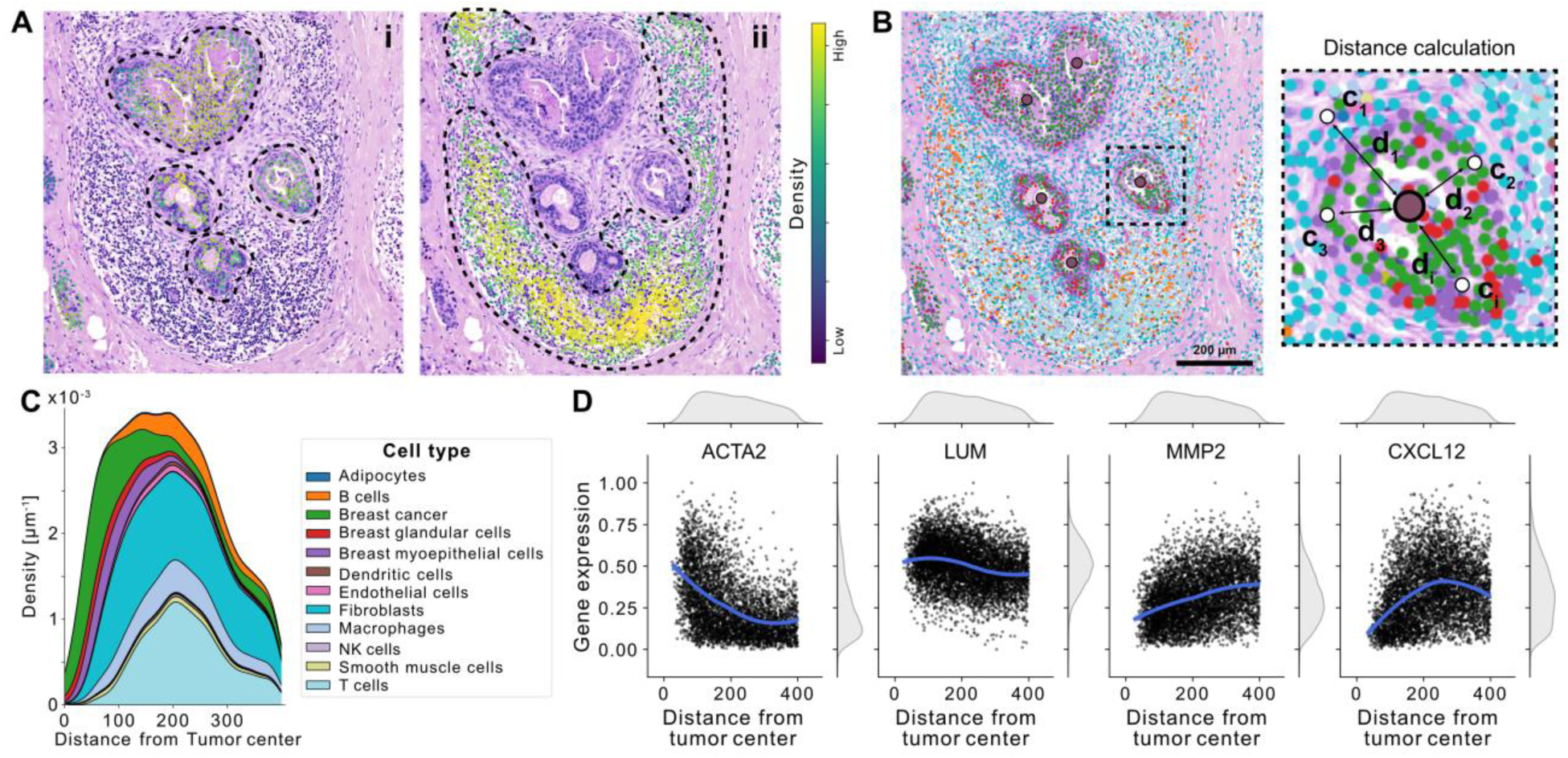
Cell type density and distance-dependent gene expression changes. **(A)** Overlay of cellular data as points and H&E stained tissue section of breast cancer example dataset. Colors denote the density of breast cancer cells (i) and T cells (ii). Density was calculated using the mellon^17^ package. **(B)** Schematic illustrating the distance calculation and annotations. In this example, a point annotation of the tumor center was used as example. **(C)** Cellular composition with increasing distance from the tumor center. Colors denote the cell types. **(D)** Scatter plot showing the gene expression of different genes with increasing distance from the tumor center. Each dot represents the gene expression in one cell. The smoothed line was calculated using LOESS regression. LOESS: Locally estimated scatterplot smoothing.

## Discussion

In this work, we introduce InSituPy, a comprehensive Python-based framework, designed to simplify the handling, analysis, and visualization of single-cell spatial transcriptomics data. While state-of-the-art Python-based analysis frameworks such as SpatialData^5^ or Squidpy^18^ focus on a sample-level analysis, InSituPy allows handling of the data on the sample level but also on an experiment level, i.e. of multiple samples in parallel. With functions to save, load, query and analyze the data on a multi-sample level, InSituPy lays the foundation for large spatial transcriptomics projects, involving e.g. large patient cohorts. For the multi-sample analysis of such datasets, InSituPy provides functionalities to conduct differential gene expression analysis within and across samples using a simple syntax. Example workflows for this analysis and others are provided in the documentation. This includes preprocessing steps such as normalization, dimensionality reduction as well as analysis steps such as GO term enrichment analysis, distance-dependent analysis and visualization of the results using different plotting functionalities. All these functionalities were demonstrated in this publication on a previously published breast cancer datasets and used to effortlessly integrated pathological knowledge into the analysis. This revealed multiple breast cancer subtypes on a single-cell level and helped to reveal differences within the subtypes between different histological regions of the dataset. The open-source publication of the framework will facilitate the implementation of additional, community-developed multi-sample analysis workflows in the future.

On a sample level, currently published frameworks such as SpatialData structure the data based on the mathematical properties of the modalities, e.g. in polygons or points. In contrast to this, InSituPy uses a data structure which groups the modalities into biologically meaningful layers, making an analysis more intuitive and more easily adaptable for non-bioinformaticians and medical experts. To improve the integration with other frameworks, functions will be implemented to convert data from the InSituPy format into the SpatialData format.

A central step in the analysis of spatial transcriptomics data is the alignment of orthogonally generated imaging data, which often requires tedious manual steps. Based on previously published analysis pipelines^19^ and similarly to tools like VoltRon^20^, InSituPy uses the computer vision toolbox OpenCV to facilitate a fast and automatic alignment of histological or immunofluorescent stainings performed subsequently to the spatial transcriptomics readout.

To verify results and develop hypotheses during analysis, a fast interactive visualization of the data is crucial. Based on the napari^8^ framework, InSituPy offers visualization of image data and cellular transcriptomic data as well as displaying and manual addition of histological annotations or regions, facilitating the integration of pathological expert knowledge and opening new ways of generating hypotheses and driving translational research forward.

While InSituPy is currently focused on the analysis of Xenium In Situ data, the data structure is suitable for all kinds of imaging-based but also sequencing-based spatial transcriptomics methodology. Future efforts will focus on expanding the set of loading functions to open InSituPy for more different methodologies. Further, we will expand the documentation and integrate more analysis functions as well as analysis workflows, focusing especially on a multi-sample analysis. We also plan to integrate InSituPy into the scverse^21^ ecosystem and use the unified data format of SpatialData for saving the data while preserving the biology-inspired data structure of InSituPy.

In conclusion, InSituPy provides an interactive analysis platform for multi-sample, single-cell spatial transcriptomics data, with a focus on histology and pathology-driven analysis, aiming to facilitate the analysis for researchers from diverse backgrounds.

## Methods

### Python package InSituPy

The InSituPy framework is built in Python v3.9 and offers an multi-sample and sample-level analysis of single-cell spatial transcriptomic data. All code is licensed under the “BSD 3-Clause” and available on Github (https://github.com/SpatialPathology/InSituPy). Information about installation and usage as well as tutorials are available on Github and additional documentation is built using Sphinx and hosted on Read the Docs.

### InSituPy framework dependencies

The InSituPy framework depends on following packages: *scanpy*, *loompy* (single-cell transcriptomics data analysis); *dask*, *opencv*, *zarr* (image data operations); *geopandas*, *rasterio* (spatial data operations); *napari* (visualization); *numpy*, *scikit*, *scipy* (general mathematical operations); *matplotlib*, *seaborn*, *adjusttext* (plotting); *pandas*, *toml, fastparquet, pyarrow* (handling of general data formats), *pytest* (code testing).

### Architecture of data classes within InSituData

To achieve optimal storage and handling of the different data modalities (images, cellular transcriptomes, cellular boundaries, transcripts, geometric data), they are structured into biologically meaningful entities using specialized data classes and collected within an *InSituData* object. Loading of a certain modality can be invoked using the respective ‘*load_{modality}*’ function (**Supplementary Table S1**). The different data classes are described subsequently. Image data is handled using the *ImageData* class, allowing the addition of multiple images and associated metadata such as resolution of the image or other OME metadata. The image resolution is pivotal because all data classes of InSituPy use metric units (usually µm) as basis for building a common coordinate system. Images can be saved either as *zarr*-formated image file^22^ or alternatively as *OME-TIFF*^23^ to allow compatibility with bioimage analysis software such as QuPath^9^. At the center of the analysis is the cellular data which is stored in a *CellData* object and consists of the single-cell transcriptomic data and the cellular boundaries of each cell. Within the CellData object the single-cell transcriptomic data is stored as *AnnData*^24^ object which allows compatibility with *scverse* analysis packages, including *SpatialData*^5^, *Squidpy*^18^ or *ScanPy*^25^. The information about cellular boundaries is stored in a *BoundariesData* object as pyramidal dask arrays allowing fast visualization in the form of a labels layer in *napari*. Information about the transcript locations and identities, as measured in in situ sequencing or in situ hybridization methods, is stored as *pandas*^26^ dataframe in the “.transcripts” attribute. The complexity of histological annotations is handled using the *ShapesData* object. Based on a unique key as reference (e.g. name of the pathologist or overarching type of the annotation), ShapesData allows the addition of geometric information. The geometric shapes are stored as shapely^27^ object inside a GeoPandas^28^ dataframe, which facilitates the application of advanced functions from both packages. Within geometric annotations, InSituPy differentiates between “annotations” and “regions”. The main difference between the two types is that an annotation is allowed to have multiple geometric objects per object name while a region is required to provide a unique name for each geometry. Annotations are meant to reflect histological annotations (e.g. tumor, necrosis) where multiple geometries with the same name are expected, while regions are meant to reflect regions such as TMA cores where each geometry possesses a unique name. These two possibilities are reflected by two daughter classes of *ShapesData*: *AnnotationsData* and *RegionsData*.

### Architecture of InSituExperiment object

To allow a comprehensive analysis of datasets with multiple samples, various *InSituData* objects can be combined in an *InSituExperiment* object, connecting the data with corresponding metadata (e.g. clinical data or experimental data). Different strategies are implemented to build an InSituExperiment object (Supp. Figure 1 A), including (i) the direct generation from an InSituData object based on regions, (ii) the addition of individual InSituData objects to an empty or existing InSituExperiment object or (iii) directly from a configuration file. Metadata can either be added using the *append_metadata* function or specified in the configuration file.

### Reading and saving of data

Both on an multi-sample and single-sample level functions are implemented to save the *InSituExperiment* or *InSituData* object to disk. After saving, the data can be read using either *InSituExperiment.read()* or *InSituData.read().* Modalities can be loaded using the respective loading functions. To load data from different single-cell spatial transcriptomics technologies, distinct reading functions are implemented (e.g. *read_xenium* for the Xenium In Situ method). Further, documentation on how to read custom data is available.

### Download of demo datasets

To facilitate a fast implementation and testing of InSituPy, functions to download multiple different demo datasets of the Xenium In Situ technology are implemented. This includes Xenium v1 datasets from different human samples: breast cancer, non-diseased kidney, pancreatic cancer, skin melanoma, brain cancer, lung cancer and lymph node. Further, a lymph node sample from the Xenium 5K technology and a test dataset with small data size can be downloaded through these functions. The URLs used for the download can be found in **Supplementary Table 3**.

### Preprocessing functions

An example preprocessing workflow for Xenium In Situ data is provided in the documentation. Important *Scanpy*-based^25^ preprocessing steps such as normalization, transformation and dimensionality reduction were summarized in the functions *normalized_and_transform* and *reduce_dimensions*, respectively. For normalization, the function uses Scanpy’s *normalize_total* function and for transformation both log-transformation and square-root transformation are implemented.

### Automated image registration of histological and immunofluorescent stainings

For the automated image registration pipeline, functions from the computer vision library OpenCV (*v4.8.0.76*) were used. This pipeline aligns the subsequently stained images to the nuclear image acquired during the scST measurement. Before the registration, both images were downscaled to a maximum width of 4000 pixels to reduce the memory consumption. *Scale-Invariant Feature Transform*^6^ was used with default parameters to identify keypoints in both images. Subsequently, the *Fast Library for Approximate Nearest Neighbors* (FLANN)^7^ was used to match the keypoints in both images using the parameters *kdtree=5* and *checks=50*. Lowe’s ratio test^29^ was used to eliminate false matches using a ratio threshold of *0.7*. If not enough matched keypoints were found, the algorithm automatically tested whether flipping the image vertically or horizontally led to better results and continues with the best of these options. Before calculation of the transformation matrix, the remaining keypoints were rescaled to the size of the original input images. For a perspective transformation, the *findHomography* function and for an affine transformation the *estimateAffine2D* function were used. In both functions, the most robust matches were selected using *random sample consensus algorithm*, followed by the calculation of an affine or perspective transformation matrix. This transformation matrix was then used to transform the histological image using the functions *warpPerspective* or *warpAffine*, respectively. A demonstration of the automated registration pipeline can be found in the documentation.

### Histological annotation in QuPath and import into InSituPy

For annotation of histological images in external software, the open-source software *QuPath*^9^ was tested and workflows for its use were implemented. The implementation of the workflows was done in version *0.5.1* of QuPath. Annotations can be applied using QuPath’s standard annotation tools and exported using *File > Export objects as GeoJSON*. To import the geometric shapes as either annotations or regions (see **Figure 1 F**), two import functions (*import_annotations* and *import_regions*) are provided. Since InSituPy uses metric units, it is important to provide a scale factor in µm per pixel. The scale factor corresponds to the resolution of the image on which the annotations were generated. To determine which cells are located within a certain annotation, the function *assign_annotations* can be used. Examples of this workflow are provided in the documentation.

### Visualization function using napari

For interactive visualization of the scST data a viewer was implemented using napari^8^ (*v0.4.18*). The viewer can be invoked from a *InSituData* or *InSituExperiment* object using the *show* function. In case of an InSituExperiment object the index of the respective dataset needs to be provided. Images are added lazily to the viewer using the *Dask* framework (*v2023.9.2*)^30^ from either a *zarr*-formatted^22^ or *OME-TIFF*^23^ formatted image file. To visualize and interact with the data, different widgets were implemented (**Supplementary Figure 3 E-N**). Cellular data can be displayed as points layer using the “Show data” widget and filtered using “Filter cells” widgets. Filtering can be used e.g. to display only a certain cell type. Cellular boundaries can be visualized as labels layer using the “Show boundaries” widget. To localize and highlight a certain cell, the “Navigate to cell” widget can be used. Annotations or regions can be added using the “Add geometries” widget. For geometric annotations or regions, a shapes layer is added while for point annotations a points layer is added to the viewer. Already existing annotations or regions can be displayed using the “Show geometries” widget.

### Differential gene expression analysis

Differential gene expression analysis is based on the *rank_genes_groups* function from Scanpy (v1.10.3). The analysis can be performed either on the sample level using the function *differential_gene_expression* or on the experiment level using the class function *dge* of an *InSituExperiment* object. Further, the analysis can be limited to annotations, regions or observation categories (e.g. cell types).

### GO term enrichment analysis

InSituPy provides functionalities to connect to APIs from different GO term enrichment analysis web servers including STRING^12^, g:profiler^13^, and Enrichr^14^. Packages used to connect to the APIs include *requests* (v2.32.3), *gprofiler-official* (v1.0.0) and *gseapy* (v1.1.4). The exact statistical tests and calculation of the false discovery rate (FDR) depend on the web server used and details can be found in the documentation of the respective publications. Enrichment scores are returned as -log10(FDR). The gene ratio was calculated as fraction of genes of a specific pathway that were present in the query list.

### Distance-dependent gene expression analysis

The function *calc_distance_of_cells_from* uses the *geopandas* package to calculate the Euclidean distance between each cell to the closest point on a specified annotation or region. If, in case of annotations, multiple annotation objects exist only the closest annotation is used as reference.

### Cellular density analysis

For cellular density analysis two different strategies are implemented to calculate the density of a specified cell type: (i) Kernel-density estimation using Gaussian kernels as implemented in *scipy* (v1.13.1), and (ii) log density estimation as implemented in *mellon*^17^ (v1.4.3). The density results of both methods can be chosen to be clipped to their 5^th^ and 95^th^ percentile as recommend in ref. ^17^.

### Plotting functions

InSituPy contains different plotting functionalities to visualize the results of above-mentioned analyses. This includes volcano plots, pie plots and dot plots which are plotted using functionalities from the packages *matplotlib* (v3.9.3), seaborn (v0.13.2), and *adjustText* (v1.3.0).

## Code availability

The Python packages InSituPy (https://github.com/SpatialPathology/InSituPy) is available on Github under the BSD-3-Clause license. Tutorials and documentation can be found in the same repository on Github.

## Acknowledgements

This work was supported by the BMBF through the SATURN3 project with the reference number 01KD2206A. Further, we want to thank the team of the Core Facility ‘Comparative Experimental Pathology’ and the tissue biobank at the Institute of Pathology of TUM for bringing in their pathological expert knowledge, especially Thomas Metzler, Tanja Groll and Leona Arps, as well as their expertise on sample preparation, especially, Aezlat Alkhamas, Olga Seelbach and Aida Mesinovic for their expertise in sample preparation. Figures and schematics were created using Affinity Designer 2. Third-party icons were retrieved from flaticon.com.

## Author contributions

J.W. conceptionalized, designed and authored the InSituPy library, with contributions from A.C., B.L., G.I., and A.E.L. J.W. wrote the manuscript. K.S. supervised the work.

## Competing interests

The authors have no competing interests to declare.

## Supplementary figures

**Supplementary Figure 1.**
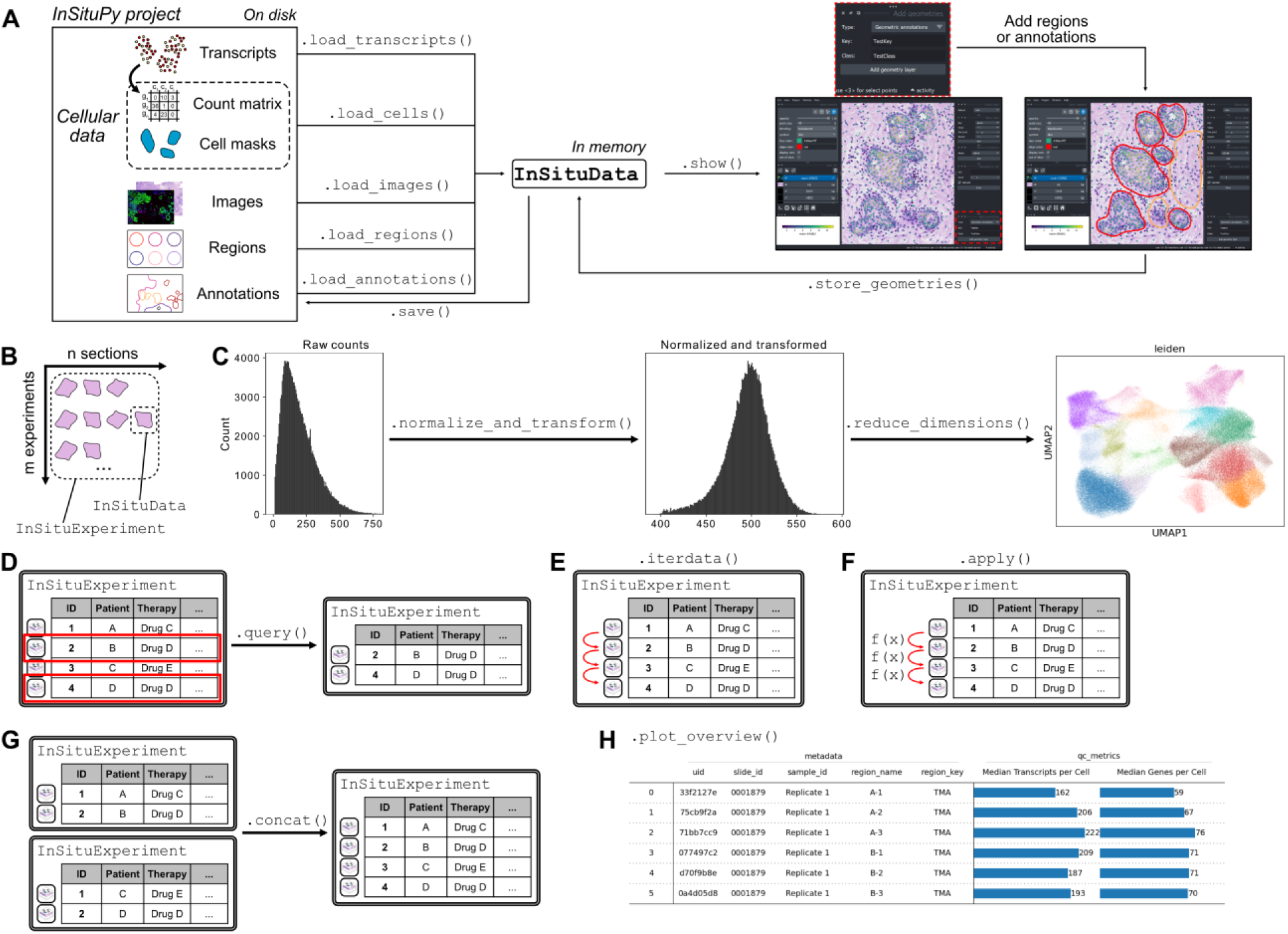
(A) Overview of functions to load the different data modalities into an InSituPy project. **(B)** Schematic showing the overarching data structure of InSituPy. The InSituData class is used to store data from single tissue sections while the InSituExperiment class collects multiple InSituData classes and integrates data across experiments. **(C)** Overview of functions for data normalization and transformation, implemented in InSituPy.

**Supplementary Figure 2.**
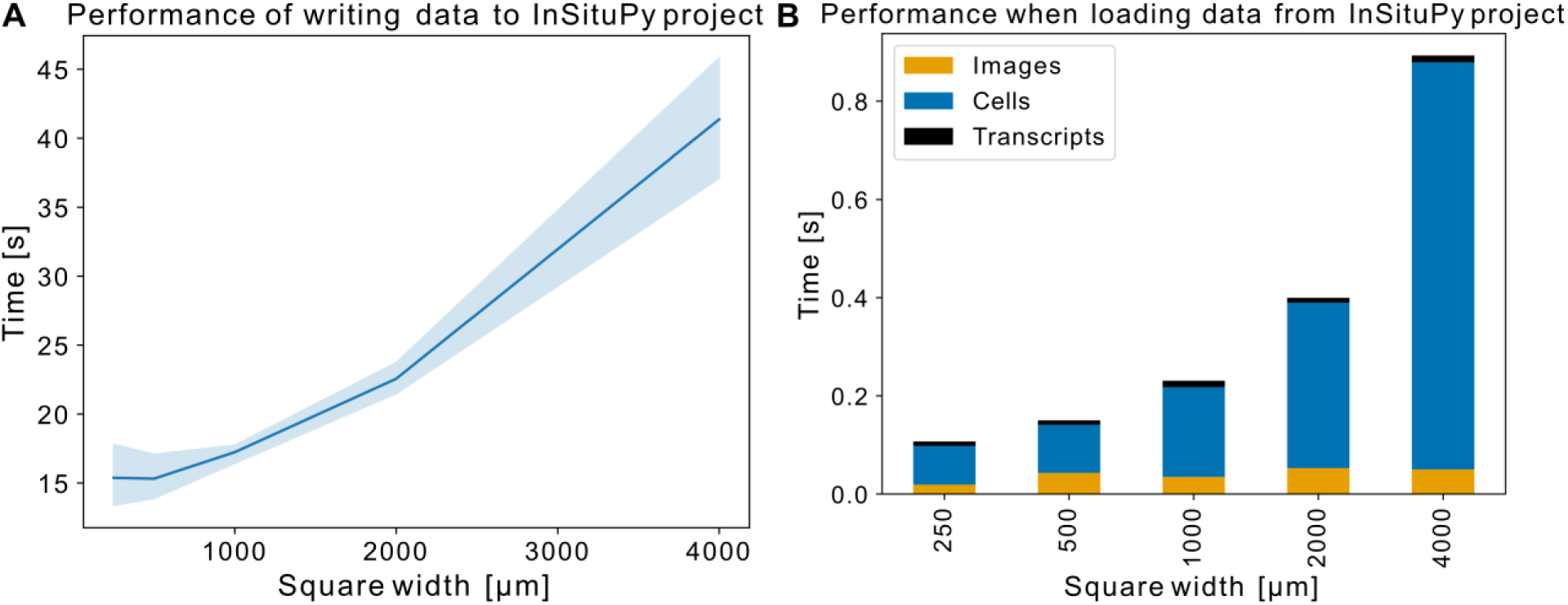
Performance tests for writing and loading data with InSituPy.

**Supplementary Figure 3.**
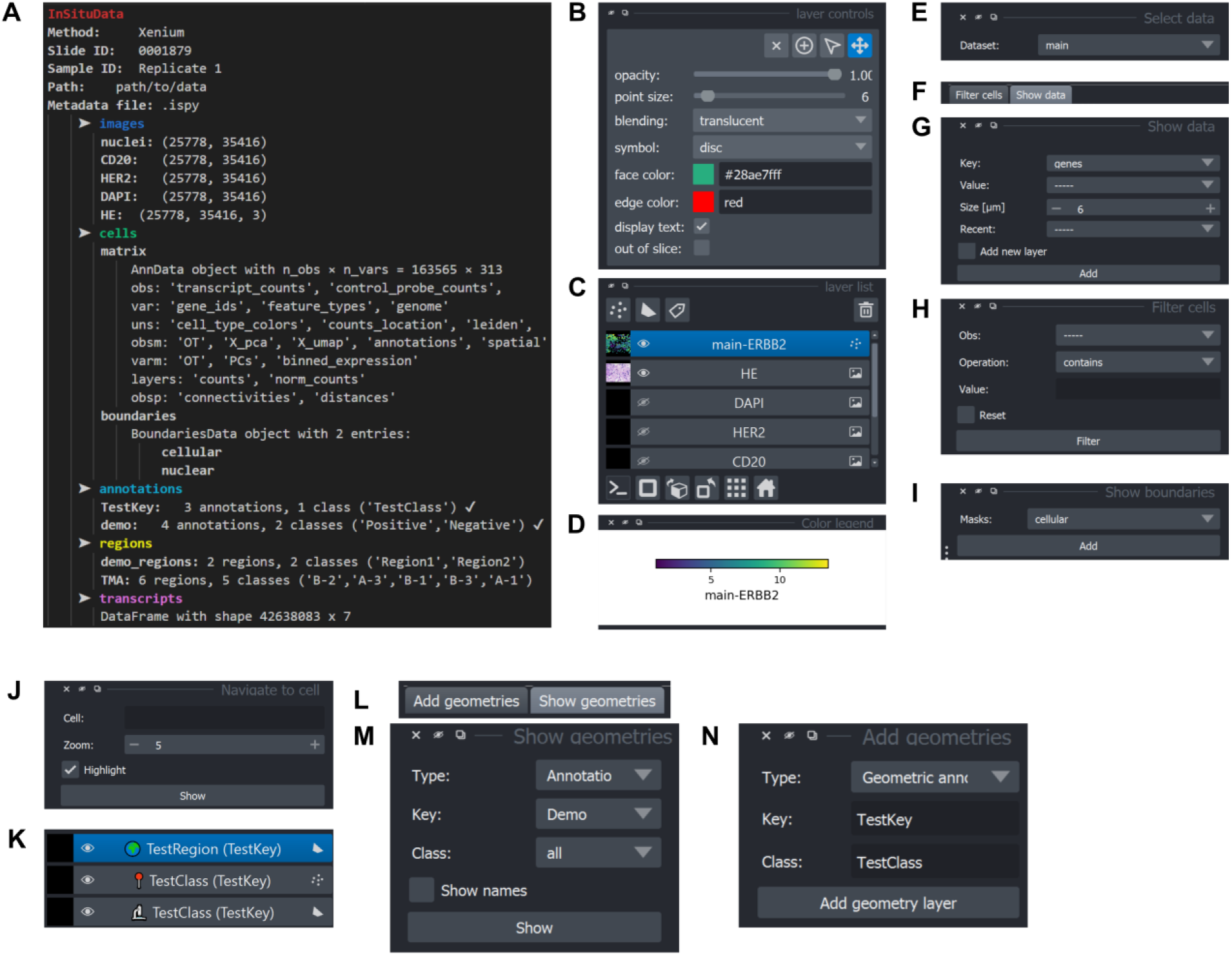
Overview of data visualizations and viewer widgets in InSituPy.

**Supplementary Figure 4.**
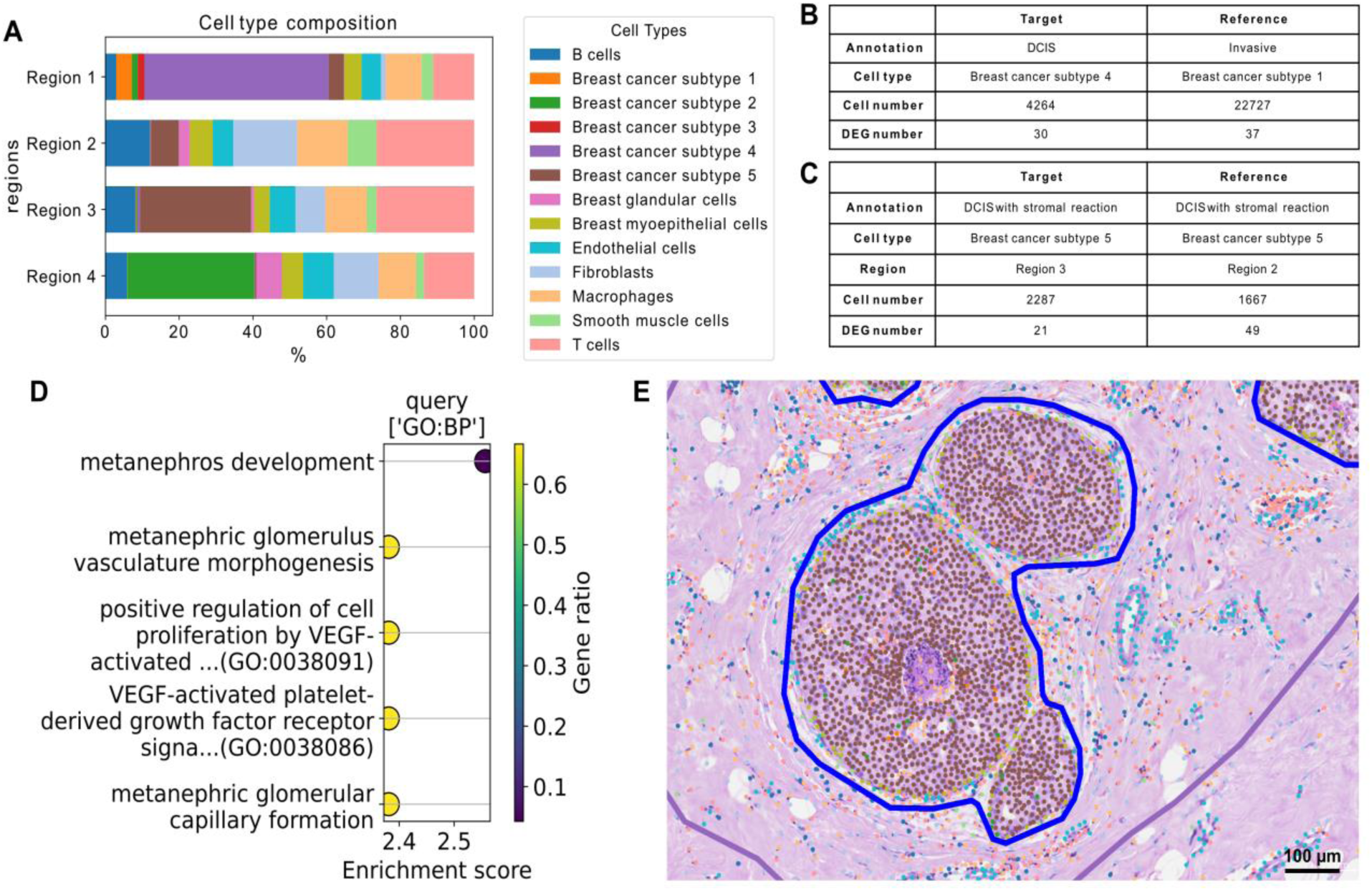
Cell composition and differential gene expression analysis with InSituPy. (A) Cell composition analysis between four selected regions in breast cancer dataset. Colors denote cell types. (B) and (C) Configuration tables corresponding to differential gene expression analyses in Figure 3F or H, respectively. (D) Dot plot showing results of gene ontology term enrichment analysis of up-regulated genes in breast cancer subtype 5 in region 3 compared to region 2. (E) Example image of breast cancer subtype 5 cells in region 3. Colors denote the cell types and correspond to the legend in A.

## Supplementary tables

**Supplementary Table 1.** Data formats used for the different data modalities in InSituPy.

| Modality | Attribute name | Loading function | Data format |  |
| --- | --- | --- | --- | --- |
|  |  |  | After loading | On disk |
| Single-cell transcriptome | .cells.matrix | load_cells() | AnnData | H5AD |
| Cellular segmentation masks | .cells.boundaries |  | Dask Array | Zarr |
| Image data | .images | load_images() | Dask Array | Zarr/OME-TIFF |
| Transcript locations and identities | .transcripts | load_transcripts() | Pandas DataFrame | Parquet |
| Histological annotations | .annotations | load_annotations() | GeoPandas DataFrame | GeoJSON |
| Regional annotations | .regions | load_regions() | GeoPandas DataFrame | GeoJSON |

**Supplementary Table 2.**
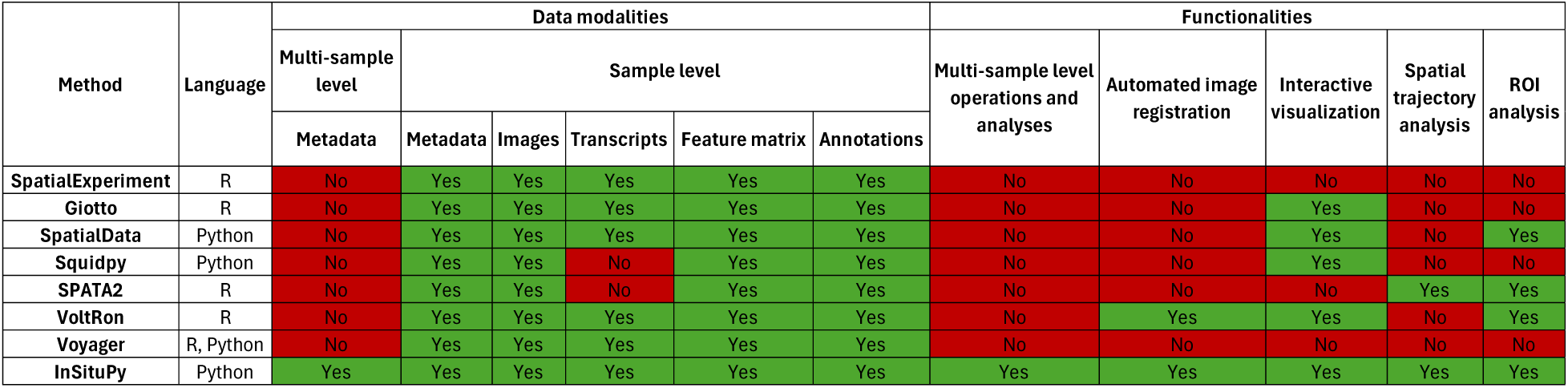
Comparison of functionalities in different frameworks for spatial transcriptomics data.

| Method | Language | Data modalities |  |  |  |  |  | Functionalities |  |  |  |  |
| --- | --- | --- | --- | --- | --- | --- | --- | --- | --- | --- | --- | --- |
|  |  | Multi-sample level | Sample level |  |  |  |  | Multi-sample level operations and analyses | Automated image registration | Interactive visualization | Spatial trajectory analysis | ROI analysis |
|  |  |  | Metadata | Images | Transcripts | Feature matrix | Annotations |  |  |  |  |  |
| SpatialExperiment | R | No | Yes | Yes | Yes | Yes | Yes | No | No | No | No | No |
| Giotto | R | No | Yes | Yes | Yes | Yes | Yes | No | No | Yes | No | No |
| SpatialData | Python | No | Yes | Yes | Yes | Yes | Yes | No | No | Yes | No | Yes |
| Squidpy | Python | No | Yes | Yes | No | Yes | Yes | No | No | Yes | No | No |
| SPATA2 | R | No | Yes | Yes | No | Yes | Yes | No | No | No | Yes | Yes |
| VoltRon | R | No | Yes | Yes | Yes | Yes | Yes | No | Yes | Yes | No | Yes |
| Voyager | R, Python | No | Yes | Yes | Yes | Yes | Yes | No | No | No | No | No |
| InSituPy | Python | Yes | Yes | Yes | Yes | Yes | Yes | Yes | Yes | Yes | Yes | Yes |

**Supplementary Table 3.**
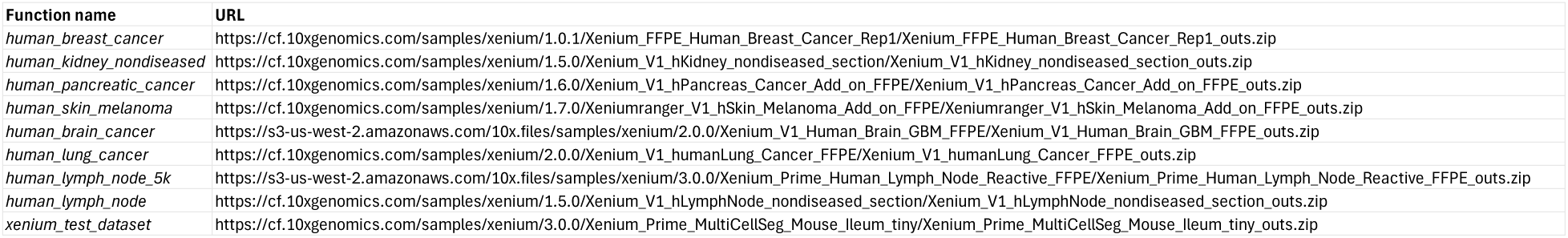
Source of example datasets implemented in InSituPy.

| Function name | URL |
| --- | --- |
| human_breast_cancer | https://cf.10xgenomics.com/samples/xenium/1.0.1/Xenium_FFPE_Human_Breast_Cancer_Rep1/Xenium_FFPE_Human_Breast_Cancer_Rep1_outs.zip |
| human_kidney_nondiseased | https://cf.10xgenomics.com/samples/xenium/1.5.0/Xenium_V1_hKidney_nondiseased_section/Xenium_V1_hKidney_nondiseased_section_outs.zip |
| human_pancreatic_cancer | https://cf.10xgenomics.com/samples/xenium/1.6.0/Xenium_V1_hPancreas_Cancer_Add_on_FFPE/Xenium_V1_hPancreas_Cancer_Add_on_FFPE_outs.zip |
| human_skin_melanoma | https://cf.10xgenomics.com/samples/xenium/1.7.0/Xeniumranger_V1_hSkin_Melanoma_Add_on_FFPE/Xeniumranger_V1_hSkin_Melanoma_Add_on_FFPE_outs.zip |
| human_brain_cancer | https://s3-us-west-2.amazonaws.com/10x.files/samples/xenium/2.0.0/Xenium_V1_Human_Brain_GBM_FFPE/Xenium_V1_Human_Brain_GBM_FFPE_outs.zip |
| human_lung_cancer | https://cf.10xgenomics.com/samples/xenium/2.0.0/Xenium_V1_humanLung_Cancer_FFPE/Xenium_V1_humanLung_Cancer_FFPE_outs.zip |
| human_lymph_node_5k | https://s3-us-west-2.amazonaws.com/10x.files/samples/xenium/3.0.0/Xenium_Prime_Human_Lymph_Node_Reactive_FFPE/Xenium_Prime_Human_Lymph_Node_Reactive_FFPE_outs.zip |
| human_lymph_node | https://cf.10xgenomics.com/samples/xenium/1.5.0/Xenium_V1_hLymphNode_nondiseased_section/Xenium_V1_hLymphNode_nondiseased_section_outs.zip |
| xenium_test_dataset | https://cf.10xgenomics.com/samples/xenium/3.0.0/Xenium_Prime_MultiCellSeg_Mouse_Ileum_tiny/Xenium_Prime_MultiCellSeg_Mouse_Ileum_tiny_outs.zip |

## Notes

https://github.com/SpatialPathology/InSituPy

### Competing Interest Statement

The authors have declared no competing interest.

## References

1. Lubeck, E., Coskun, A. F., Zhiyentayev, T., Ahmad, M. & Cai, L. Single-cell in situ RNA profiling by sequential hybridization. Nat. Methods 11, 360–361 (2014).

2. Xia, C., Fan, J., Emanuel, G., Hao, J. & Zhuang, X. Spatial transcriptome profiling by MERFISH reveals subcellular RNA compartmentalization and cell cycle-dependent gene expression. Proc. Natl. Acad. Sci. 116, 19490–19499 (2019).

3. Ke, R. et al. In situ sequencing for RNA analysis in preserved tissue and cells. Nat. Methods 10, 857 (2013).

4. Lee, J. H. et al. Highly Multiplexed Subcellular RNA Sequencing in Situ. Science 343, 1360–1363 (2014).

5. Marconato, L. et al. SpatialData: an open and universal data framework for spatial omics. Nat. Methods 1–5 (2024) doi:10.1038/s41592-024-02212-x.

6. Lowe, D. G. Distinctive Image Features from Scale-Invariant Keypoints. Int. J. Comput. Vis. 60, 91–110 (2004).

7. Muja, M. & Lowe, D. G. Fast Approximate Nearest Neighbors with Automatic Algorithm Configuration. in International Conference on Computer Vision Theory and Applications (2009).

8. Sofroniew, N., et al. napari: a multi-dimensional image viewer for Python. Zenodo 10.5281/zenodo.13863809 (2024).

9. Bankhead, P. et al. QuPath: Open source software for digital pathology image analysis. Sci. Rep. 7, 16878 (2017).

10. Janesick, A. et al. High resolution mapping of the tumor microenvironment using integrated single-cell, spatial and in situ analysis. Nat. Commun. 14, 8353 (2023).

11. 10X Genomics. High resolution mapping of the breast cancer tumor microenvironment using integrated single cell, spatial and in situ analysis of FFPE tissue.

12. Szklarczyk, D. et al. STRING v11: protein–protein association networks with increased coverage, supporting functional discovery in genome-wide experimental datasets. Nucleic Acids Res. 47, D607–D613 (2019).

13. Raudvere, U. et al. g:Profiler: a web server for functional enrichment analysis and conversions of gene lists (2019 update). Nucleic Acids Res. 47, W191–W198 (2019).

14. Kuleshov, M. V. et al. Enrichr: a comprehensive gene set enrichment analysis web server 2016 update. Nucleic Acids Res. 44, W90–W97 (2016).

15. The Gene Ontology Consortium. The Gene Ontology resource: enriching a GOld mine. Nucleic Acids Res (2021) doi:10.1093/nar/gkaa1113.

16. Salas, S. M. et al. Optimizing Xenium In Situ data utility by quality assessment and best practice analysis workflows. 2023.02.13.528102 Preprint at 10.1101/2023.02.13.528102 (2023).

17. Otto, D. J., Jordan, C., Dury, B., Dien, C. & Setty, M. Quantifying cell-state densities in single-cell phenotypic landscapes using Mellon. Nat. Methods 21, 1185–1195 (2024).

18. Palla, G. et al. Squidpy: a scalable framework for spatial omics analysis. Nat. Methods 19, 171–178 (2022).

19. Wirth, J. et al. Spatial transcriptomics using multiplexed deterministic barcoding in tissue. Nat. Commun. 14, 1523 (2023).

20. Manukyan, A. et al. VoltRon: A Spatial Omics Analysis Platform for Multi-Resolution and Multi-omics Integration using Image Registration. 2023.12.15.571667 Preprint at 10.1101/2023.12.15.571667 (2023).

21. Virshup, I. et al. The scverse project provides a computational ecosystem for single-cell omics data analysis. Nat. Biotechnol. 41, 604–606 (2023).

22. Miles, A., et al. zarr-developers/zarr-python: v2.4.0. Zenodo 10.5281/zenodo.3773450 (2020).

23. Besson, S. et al. Bringing Open Data to Whole Slide Imaging. Digit. Pathol. 15th Eur. Congr. ECDP 2019 Warwick UK April 10-13 2019 Proc. Eur. Congr. Digit. Pathol. 15th 2019 Warwick Engl. 2019, 3–10 (2019).

24. Virshup, I., Rybakov, S., Theis, F. J., Angerer, P. & Wolf, F. A. anndata: Annotated data. *bioRxiv* 2021.12.16.473007 (2021) doi:10.1101/2021.12.16.473007.

25. Wolf, F. A., Angerer, P. & Theis, F. J. SCANPY: Large-scale single-cell gene expression data analysis. Genome Biol. 19, 15 (2018).

26. Mckinney, W. Data Structures for Statistical Computing in Python. Proc. 9th Python Sci. Conf. (2010).

27. Gillies, S. & others. Shapely: manipulation and analysis of geometric objects. (2007).

28. Jordahl, K. GeoPandas: Python tools for geographic data. URL Httpsgithub Comgeopandasgeopandas (2014).

29. Lowe, D. G. Distinctive Image Features from Scale-Invariant Keypoints. Int. J. Comput. Vis. 60, 91–110 (2004).

30. Rocklin, M. Dask: Parallel Computation with Blocked algorithms and Task Scheduling. in Proceedings of the 14th Python in Science Conference (eds. Huff, K. & Bergstra, J.) 130–136 (2015).

